# The molecular basis of extensively drug-resistant *Salmonella* Typhi isolates from pediatric septicemia patients

**DOI:** 10.1101/2021.06.20.449163

**Authors:** Chanmi Kim, Iqra Latif, Durga P. Neupane, Gi Young Lee, Ryan S. Kwon, Alia Batool, Qasim Ahmed, Muhammad Usman Qamar, Jeongmin Song

## Abstract

Sepsis is a syndromic response to infections and is becoming an emerging threat to the public health sector, particularly in developing countries. *Salmonella* Typhi (*S*. Typhi), the cause of typhoid fever, is one primary cause of pediatric sepsis in typhoid endemic areas. Extensively drug-resistant (XDR) *S*. Typhi is more common among pediatric patients, which is responsible for over 90% of the reported XDR typhoid cases, but the majority of antibiotic resistance studies available have been carried out using *S*. Typhi isolates from adult patients. Here, we characterized XDR *S*. Typhi isolates from a medium size cohort of pediatric typhoid patients to determine their antibiotic-resistance-related gene signatures associated with common treatment options to typhoid fever patients. This study informs the molecular basis of antibiotic-resistance among recent *S*. Typhi isolates from pediatric septicemia patients, therefore providing insights into the development of molecular detection methods and control strategies for XDR *S*. Typhi.

## Introduction

Sepsis is a syndromic response to infections and is becoming an emerging threat to the public health sector. The World Health Organization (WHO) estimated that 48.9 million cases of sepsis have been reported and that one person dies every 2.58 seconds around the world (1). Furthermore, 20 million cases were detected among children, with 2.9 million deaths worldwide and 85% of these deaths in developing countries (1). *Salmonella enterica* serovar Typhi (*S*. Typhi), the causative agent of typhoid fever, is one primary cause of pediatric sepsis in typhoid endemic areas including Pakistan (1-3). Antibiotics are the primary treatment options for typhoid fever, but *Salmonella* are continuously evolving to acquire plasmid, prophage, transposon, or chromosomal gene mutation to attain resistance against antibiotics. A myriad of reports indicated the global spread of *S*. Typhi that is resistant to all of the first-line antibiotics, ampicillin, chloramphenicol, and co-trimoxazole, collectively known as multidrug-resistant (MDR) (4-8). All of the identified MDR *S*. Typhi carry the IncHI1 region located on either the plasmid or chromosome, which encodes several antibiotic-resistance genes, including *catA1* (conferring resistance to chloramphenicol), *blaTEM-1* (resistance to ampicillin), *dhfR7*, and *sul1* (resistance to trimethoprim-sulfamethoxazole), among other antibiotics-resistance-related genes found in MDR *S*. Typhi (4, 9-11).

Fluoroquinolones were used to treat MDR cases but became largely ineffective in some endemic regions. Fluoroquinolone-resistant *S*. Typhi encodes the quinolone resistance gene *qnrS* and point mutations in the quinolone resistance determining region (QRDR) harboring the genes for gyrase/topoisomerase II *gyrA* and *gyrB* and topoisomerase IV *parC* and *parE*. For instance, several point mutations occurred in *gyrA* have been correlated to resistance to fluoroquinolones, including M52L, G81C, D82G, S83F/Y/L, D87N/G/A/Y/H, and A119E (10, 12-16). Point mutations in *gyrB, parC*, and *parE* have also been reported, while some variants have been reported only from certain geographical locations (13, 15-20). Some of those mutation sites are near the quinolone binding site, which in many cases results in the inhibition of the binding of antibiotics to topoisomerases (21). *S*. Typhi strains resistant to chloramphenicol, ampicillin, co-trimoxazole, fluoroquinolones, and third-generation cephalosporins were first reported in Hyderabad, Sindh, Pakistan, affecting over 300 cases in 2016 (9), collectively known as extensively drug-resistant (XDR) *S*. Typhi (8, 22). XDR *S*. Typhi isolates commonly harbor an IncY plasmid carrying the extended-spectrum ◸-lactamase resistance gene *blaCTX-M-15* and quinolone resistance gene *qnrS*, among others (9).

Drug efflux pump systems also play a significant role in resistance to a wide range of antibiotics. *Salmonella spp*. possess five efflux pump families, including the ATP-binding cassette (ABC) MacAB-TolC system and resistance-nodulation-cell division (RND) AcrAB-TolC system (23). Members of the other families of drug transporters, major facilitator superfamily (MFS), multidrug and toxin extrusion (MATE), and small multidrug resistance (SMR), are located in the inner membrane (IM) of gram-negative bacteria (24). They usually function as independent units in the IM to translocate antibiotics across the membrane bilayer, followed by their cooperation with RND-type efflux pumps to pump out antibiotics across the entire cell envelope (24). In typhoidal *Salmonella*, point mutations at amino acid position 717 (R717Q or R717L) on AcrB, the antibiotic-binding subunit of the RND-type AcrAB-TolC efflux pump, have been correlated with resistance to azithromycin in *S*. Typhi and *S*. Paratyphi A, respectively (19, 25, 26).

Antibiotic-resistant *S*. Typhi infection is more common among children; more than 90% of the XDR typhoid cases are currently from children younger than 15 years old of age (9, 27, 28). The typhoid mortality in the pre-antibiotic era was approximately 25% (29). Typhoid fever is only partly preventable by vaccines, and two types of typhoid fever vaccines, the live-attenuated Ty21a and Vi and its conjugate subunit vaccines, are currently available (30). These vaccines exhibit the efficacy of approximately 55-85%, with the Vi-protein conjugate subunit vaccine being the most efficacious (30-32). The recent Vi-protein conjugate subunit vaccine has been demonstrated efficacious among children who are older than six months of age. There are no vaccines for early-life populations younger than six months available (31). Furthermore, *S*. Typhi strains that lack Vi have emerged in some endemic areas (33), which is most likely to make current subunit vaccines ineffective against those variants.

Macrolides (e.g., azithromycin) and carbapenems (e.g., imipenem, meropenem) remain to be “last resort” oral and injectable antibiotics for treating *S*. Typhi infection, respectively. *S*. Typhi strains resistant to macrolide azithromycin have been emerged (34). *S*. Typhi strains resistant to carbapenem antibiotic meropenem, have also been reported, and many cases of invasive nontyphoidal Salmonellae (NTS) resistant to carbapenems have been reported (25, 35-37). Given that typhoid fever vaccines and treatment options have limitations, there is an urgent need for closely monitoring drug-resistance profiles of *S*. Typhi strains at the point-of-care to provide valuable insights into the development of control strategies against drug-resistant *S*. Typhi.

## Results

### XDR *S*. Typhi strains isolated from children at various developmental stages

We isolated *S*. Typhi strains from 45 typhoid fever-suspected pediatric septicemia patients with 1-13 years of age (68.89% male and 31.11% female) who have visited the Fatima Memorial Hospital, Lahore, Punjab, Pakistan, between October 2019 to January 2020. Punjab is the most populous province approximately 1,044 km away from Hyderabad, Sindh, Pakistan, where the first XDR *S*. Typhi was reported (Figure 1A). Before samples were transferred to researchers, all *S*. Typhi samples were de-identified, number-based identification codes were assigned to samples (Table 1). The data were analyzed anonymously throughout the study. Positive blood culture bottles from the initial step using a fully automated culture and test system for patient blood specimens were sub-cultured on blood and MacConkey agar plates and incubated overnight at 37°C. Preliminary identification of the isolates was conducted according to colony morphology and culture characteristics, followed by polymerase chain reaction (PCR) and Sanger sequencing-based molecular determination (Figure 1B and C). These results indicate that all suspected patient specimens carried *S*. Typhi and all isolates are XDR since they are resistant to both first- and second-line antibiotics (Figure 1 and Table 1). XDR *S*. Typhi isolates exhibit similar minimum inhibitory concentration (MIC) values across 45 isolates with some variations, indicating that antibiotic resistance phenotypes do not correlate with age, sex, or hospital wards (Table 1).

**Figure 1.**
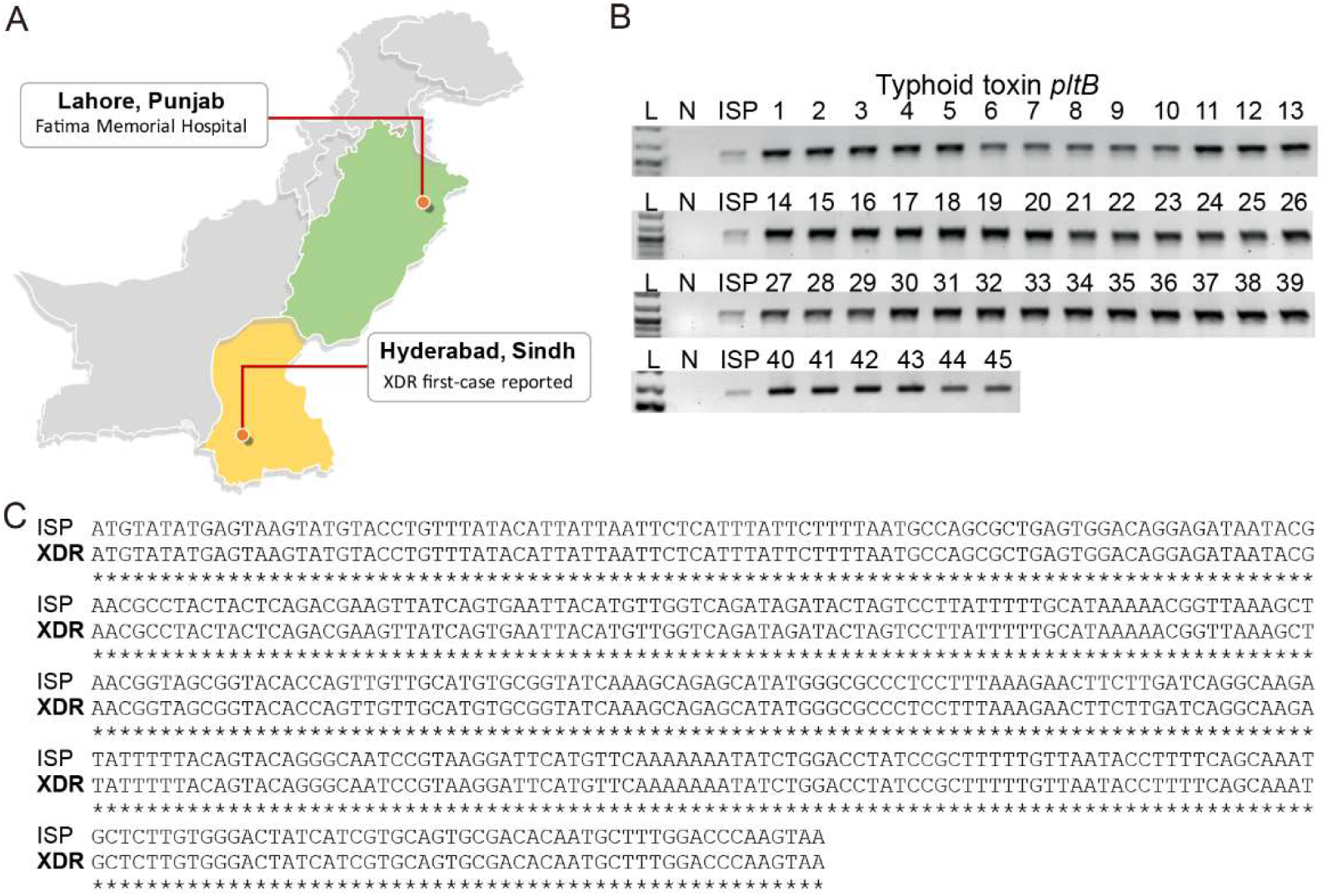
XDR *S*. Typhi strains isolated from children ages 1-13 years old. **A**, Cartoon showing the geographical location that XDR *S*. Typhi characterized in this study was isolated compared to the location that the first case of XDR *S*. Typhi was reported. **B**, PCR reactions of 45 XDR isolates for typhoid toxin *pltB*. **C**, *pltB* sequences from all 45 XDR isolates (XDR, one representative is shown) were identical to typhoid toxin *pltB* sequence of *S*. Typhi ISP2825 (ISP). ISP, antibiotic-susceptible *S*. Typhi ISP2825. N, negative control containing all PCR components except for *S*. Typhi genomic DNA. See Table 1 for sample information.

**Table 1.** The minimum inhibitory concentration (MIC) results (µg/ml) of all samples used in this study, related to Fig. 1.

| # | Age<br>* | Sex | Wards<br>** | AMP<br>*** | SXT | CIP | CTX | CRO | PIP/TZB | AMC | AZM | IPM | MEM |
| --- | --- | --- | --- | --- | --- | --- | --- | --- | --- | --- | --- | --- | --- |
| 1 | 5 | M | E | ≥32 | ≥4/76 | ≥64 | 8 | 32 | 32/4 - 64/4 | ≥32/16 | 8 | 0.25 | 0.25 |
| 2 | 2 | M | I | ≥32 | ≥4/76 | 32 | 8 | ≥64 | 4/4-128/4 | 8/4 | 2 | 0.5 | 0.25 |
| 3 | 4 | F | E | ≥32 | ≥4/76 | ≥64 | ≥64 | 32 | 32/4 - 64/4 | 16/8 | 8 | 0.5 | 0.25 |
| 4 | 10 | M | I | ≥32 | ≥4/76 | 32 | 32 | 8 | 32/4 - 64/4 | 16/8 | 4 | 0.5 | 0.25 |
| 5 | 11 | M | M | ≥32 | ≥4/76 | 8 | ≥64 | ≥64 | 32/4 - 64/4 | ≥32/16 | 4 | 0.25 | 0.5 |
| 6 | 11 | M | C | ≥32 | ≥4/76 | ≥64 | 32 | ≥64 | 4/4-128/4 | 8/4 | 4 | 0.25 | 0.5 |
| 7 | 13 | M | E | ≥32 | ≥4/76 | ≥64 | ≥64 | 32 | 4/4-128/4 | 4/2 | 4 | 0.25 | 0.5 |
| 8 | 11 | M | E | ≥32 | ≥4/76 | 32 | ≥64 | ≥64 | ≥128/4 | ≥32/16 | 4 | 0.5 | 0.5 |
| 9 | 3 | M | C | ≥32 | ≥4/76 | 8 | ≥64 | ≥64 | 4/4-128/4 | 4/2 | 4 | 0.25 | 0.5 |
| 10 | 4 | M | E | ≥32 | ≥4/76 | ≥64 | 32 | ≥64 | ≥128/4 | ≥32/16 | 4 | 0.5 | 0.5 |
| 11 | 4 | F | G | ≥32 | ≥4/76 | ≥64 | 32 | 32 | ≥128/4 | ≥32/16 | 2 | 0.5 | 0.5 |
| 12 | 4 | F | E | ≥32 | ≥4/76 | ≥64 | 8 | ≥64 | 32/4 - 64/4 | 16/8 | 2 | 0.5 | 0.25 |
| 13 | 5 | M | O | ≥32 | ≥4/76 | 32 | ≥64 | 32 | ≥128/4 | 16/8 | 2 | 0.5 | 0.5 |
| 14 | 4 | M | I | ≥32 | ≥4/76 | ≥64 | ≥64 | ≥64 | ≥128/4 | 16/8 | 8 | 0.5 | 0.5 |
| 15 | 9 | F | O | ≥32 | ≥4/76 | 32 | 32 | ≥64 | 4/4-128/4 | 8/4 | 2 | 0.5 | 0.5 |
| 16 | 2 | F | O | ≥32 | ≥4/76 | ≥64 | ≥64 | ≥64 | ≥128/4 | 16/8 | 8 | 0.5 | 0.25 |
| 17 | 3 | M | E | ≥32 | ≥4/76 | ≥64 | ≥64 | 32 | 32/4 - 64/4 | ≥32/16 | 2 | 0.25 | 0.25 |
| 18 | 6 | M | C | ≥32 | ≥4/76 | ≥64 | ≥64 | 32 | 32/4 - 64/4 | 16/8 | 8 | 0.25 | 0.25 |
| 19 | 8 | M | C | ≥32 | ≥4/76 | ≥64 | ≥64 | ≥64 | ≥128/4 | 16/8 | 8 | 0.25 | 0.25 |
| 20 | 2 | M | W | ≥32 | ≥4/76 | 32 | ≥64 | 32 | 32/4 - 64/4 | 16/8 | 8 | 0.25 | 0.25 |
| 21 | 6 | M | W | ≥32 | ≥4/76 | ≥64 | 32 | ≥64 | ≥128/4 | ≥32/16 | 2 | 0.25 | 0.25 |
| 22 | 5 | M | E | ≥32 | ≥4/76 | ≥64 | ≥64 | 32 | 32/4 - 64/4 | ≥32/16 | 8 | 0.5 | 0.25 |
| 23 | 2 | M | I | ≥32 | ≥4/76 | ≥64 | ≥64 | ≥64 | 32/4 - 64/4 | 16/8 | 4 | 0.5 | 0.5 |
| 24 | 7 | M | MI | ≥32 | ≥4/76 | ≥64 | 8 | 32 | 32/4 - 64/4 | ≥32/16 | 8 | 0.5 | 0.5 |
| 25 | 5 | M | E | ≥32 | ≥4/76 | 8 | ≥64 | ≥64 | ≥128/4 | 16/8 | 4 | 0.25 | 0.25 |
| 26 | 11 | M | E | ≥32 | ≥4/76 | ≥64 | ≥64 | 32 | ≥128/4 | 16/8 | 8 | 0.25 | 0.25 |
| 27 | 2 | M | E | ≥32 | ≥4/76 | ≥64 | ≥64 | 32 | ≥128/4 | ≥32/16 | 8 | 0.25 | 0.5 |
| 28 | 7 | M | W | ≥32 | ≥4/76 | ≥64 | 32 | ≥64 | ≥128/4 | 16/8 | 4 | 0.5 | 0.5 |
| 29 | 2 | F | W | ≥32 | ≥4/76 | ≥64 | 32 | 8 | 4/4-128/4 | 8/4 | 4 | 0.5 | 0.25 |
| 30 | 6 | M | E | ≥32 | ≥4/76 | ≥64 | ≥64 | ≥64 | 32/4 - 64/4 | 16/8 | 4 | 0.5 | 0.25 |
| 31 | 3 | M | C | ≥32 | ≥4/76 | ≥64 | ≥64 | 8 | 32/4 - 64/4 | ≥32/16 | 8 | 0.25 | 0.25 |
| 32 | 5 | M | E | ≥32 | ≥4/76 | ≥64 | 32 | 32 | 32/4 - 64/4 | 16/8 | 4 | 0.25 | 0.25 |
| 33 | 3 | F | E | ≥32 | ≥4/76 | ≥64 | ≥64 | ≥64 | 32/4 - 64/4 | 16/8 | 4 | 0.25 | 0.25 |
| 34 | 6 | M | E | ≥32 | ≥4/76 | 32 | ≥64 | ≥64 | 32/4 - 64/4 | ≥32/16 | 4 | 0.5 | 0.5 |
| 35 | 6 | F | O | ≥32 | ≥4/76 | ≥64 | ≥64 | ≥64 | 32/4 - 64/4 | 8/4 | 4 | 0.25 | 0.5 |
| 36 | 11 | M | C | ≥32 | ≥4/76 | ≥64 | ≥64 | 8 | ≥128/4 | 8/4 | 8 | 0.25 | 0.5 |
| 37 | 2 | F | W | ≥32 | ≥4/76 | 32 | 32 | ≥64 | 4/4-128/4 | 8/4 | 4 | 0.5 | 0.5 |
| 38 | 13 | F | F | ≥32 | ≥4/76 | 32 | ≥64 | ≥64 | 32/4 - 64/4 | 8/4 | 4 | 0.5 | 0.5 |
| 39 | 10 | F | E | ≥32 | ≥4/76 | ≥64 | ≥64 | 8 | 32/4 - 64/4 | 8/4 | 8 | 0.5 | 0.25 |
| 40 | 4 | F | O | ≥32 | ≥4/76 | 8 | ≥64 | ≥64 | 32/4 - 64/4 | 8/4 | 4 | 0.25 | 0.25 |
| 41 | 5 | M | O | ≥32 | ≥4/76 | ≥64 | ≥64 | 32 | ≥128/4 | ≥32/16 | 8 | 0.25 | 0.5 |
| 42 | 5 | M | E | ≥32 | ≥4/76 | ≥64 | 32 | ≥64 | 32/4 - 64/4 | 16/8 | 4 | 0.25 | 0.25 |
| 43 | 8 | F | E | ≥32 | ≥4/76 | 32 | 32 | ≥64 | ≥128/4 | 8/4 | 4 | 0.25 | 0.5 |
| 44 | 2 | F | O | ≥32 | ≥4/76 | ≥64 | ≥64 | ≥64 | 32/4 - 64/4 | ≥32/16 | 4 | 0.5 | 0.25 |
| 45 | 1 | M | E | ≥32 | ≥4/76 | ≥64 | 8 | 32 | 32/4 - 64/4 | ≥32/16 | 4 | 0.25 | 0.25 |

\*, Age in years.
\*\*, E, Peads Emergency; I, PAED-ICU; M, Male Medical; C, Clinical Laboratory; G, General OPD; O, Peads Medical OPD; W, Pead Medical Ward; MI, Pead Medical ICU; F, FMH-executive clinic.
\*\*\*, Antibiotic acronym: AMP, ampicillin, SXT, trimethoprim-sulfamethoxazole, CIP, ciprofloxacin, CTX, cefotaxime, CRO, ceftriaxone, AZM, azithromycin, PIP, piperacillin, TZB, tazobactam, AMC, amoxicillin/ clavulanic acid, IPM, imipenem, MEM, meropenem.

### Molecular basis of resistance to first-line antibiotics among XDR *S*. Typhi isolates

We hypothesized that resistance to first- and second-line antibiotics among XDR *S*. Typhi isolates is primarily due to the acquisition and/mutation of antibiotic-resistance-related genes. To carry out a series of molecular characterization via PCR and/or PCR amplicon sequencing, we selected 18 *S*. Typhi samples based on sex, age, hospital wards, antibiotic-resistance profile, and MIC (Table 2). According to child development milestones defined by the Centers for Disease Control and Prevention (CDC), age groups were split into toddlers (ages 1 to 2), preschoolers (ages 3 to 5), school-age children (ages 6 to 12), and adolescents (ages 13 to 18) (Figure 2A). To understand the molecular basis of the MDR phenotype resistant to all the first-line antibiotics with clinical relevance, we have designed primers specific to *catA1, blaTEM1, dhfR7*, and *sul1*, MDR-related genes encoded in the IncHI1 region (Table 3). Using these primers, we evaluated the presence of these MDR-related genes in 18 selected *S*. Typhi isolates via PCR analysis. All PCR reactions resulted in amplicons with expected size for specific genes except for two controls, antibiotic-susceptible *S*. Typhi ISP2825 (ISP) and a control PCR reaction mixture that did not contain any *S*. Typhi genomic DNA (N) (Figure 2B-D). Drug-susceptible *S*. Typhi ISP clinical isolate available in the laboratory was used as a control since all *S*. Typhi isolates from this cohort were XDR (Table 1). These results indicate that the MDR phenotype exhibited by these XDR *S*. Typhi isolates is most likely due to MDR-related genes encoded in the IncHI1 region.

**Table 2.** Select of samples for molecular characterization, related to Figs 2-6.

| Lab No | Age (Yr) | Sex | AMP | SXT | CIP | CTX | CRO | PIP/TZB | AMC/CLA | AZM | IPM | MEM |
| --- | --- | --- | --- | --- | --- | --- | --- | --- | --- | --- | --- | --- |
|  |  |  | MDR |  | XDR |  |  |  |  |  |  |  |
| Toddlers (1-2 yrs) |  |  |  |  |  |  |  |  |  |  |  |  |
| 16 | 2 | F | R | R | R | R | R | I | I | S | S | S |
| 2 | 2 | M | R | R | R | R | R | S | S | S | S | S |
| 20 | 2 | M | R | R | R | R | R | I | I | S | S | S |
| 23 | 2 | M | R | R | R | R | R | I | I | S | S | S |
| 27 | 2 | M | R | R | R | R | R | I | I | S | S | S |
| Pre-schoolers (3-5 yrs) |  |  |  |  |  |  |  |  |  |  |  |  |
| 33 | 3 | F | R | R | R | R | R | I | I | S | S | S |
| 9 | 3 | M | R | R | R | R | R | S | S | S | S | S |
| 17 | 3 | M | R | R | R | R | R | I | I | S | S | S |
| 31 | 3 | M | R | R | R | R | R | I | I | S | S | S |
| 40 | 4 | F | R | R | R | R | R | I | S | S | S | S |
| 12 | 4 | F | R | R | R | R | R | I | I | S | S | S |
| 1 | 5 | M | R | R | R | R | R | I | I | S | S | S |
| School-age children (6-12 yrs) |  |  |  |  |  |  |  |  |  |  |  |  |
| 15 | 9 | F | R | R | R | R | R | S | S | S | S | S |
| 6 | 11 | M | R | R | R | R | R | S | S | S | S | S |
| 8 | 11 | M | R | R | R | R | R | I | I | S | S | S |
| 36 | 11 | M | R | R | R | R | R | I | S | S | S | S |
| Adolescents (13-18 yrs) |  |  |  |  |  |  |  |  |  |  |  |  |
| 38 | 13 | F | R | R | R | R | R | I | S | S | S | S |
| 7 | 13 | M | R | R | R | R | R | S | S | S | S | S |

**Table 3.**
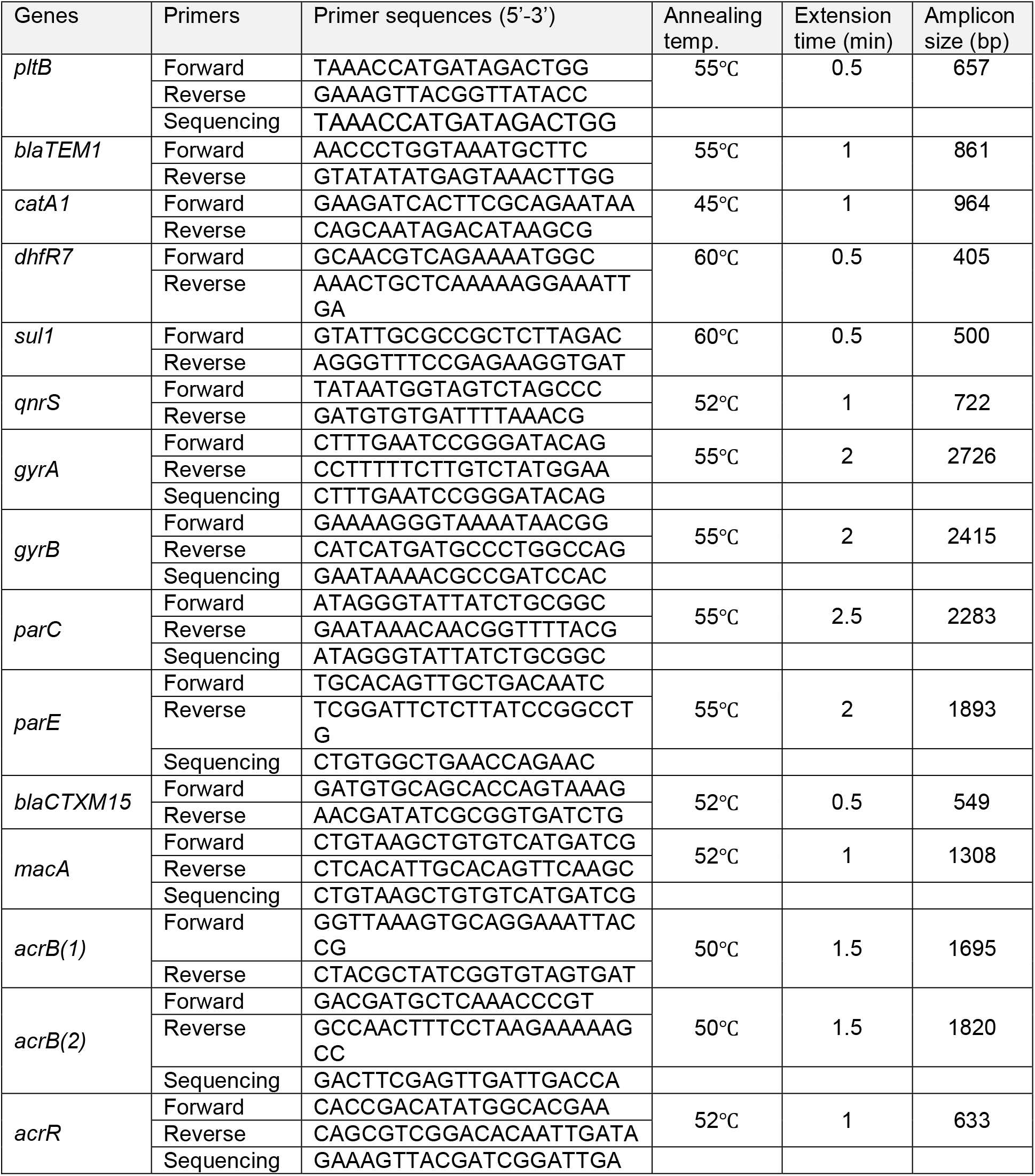
PCR and sequencing primers and PCR conditions used in this study, related to Methods.

**Figure 2.**
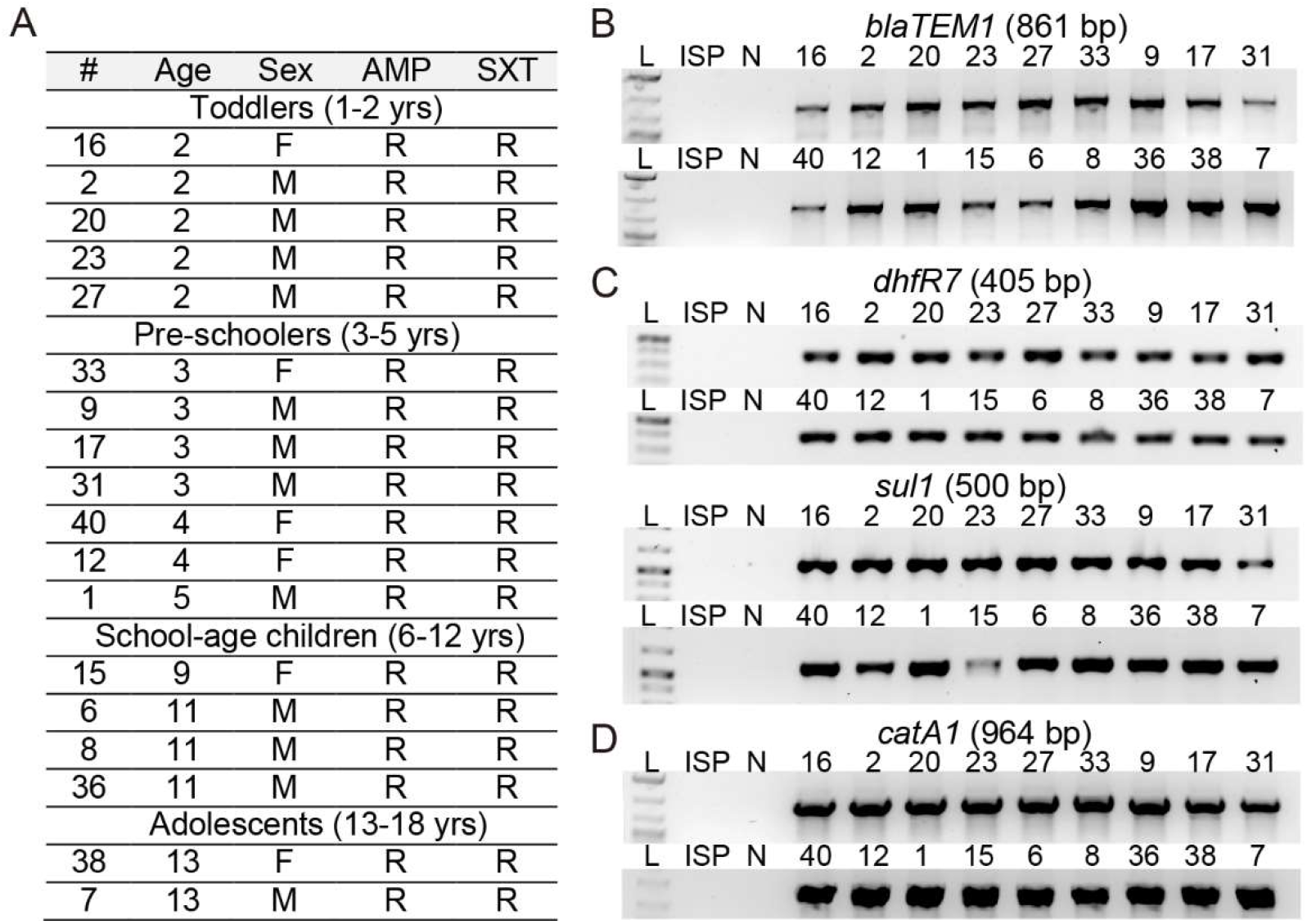
Molecular basis of resistance to first-line antibiotics among XDR *S*. Typhi isolates. **A**, Sample information and antibiotic resistance profiles of select of XDR *S*. Typhi isolates for molecular characterization. See Table 2 for details. **B**, PCR reactions for *blaTEM1*. **C**, PCR reactions for *dhfR7* and *sul1*. **D**, PCR reactions for *catA1*. ISP, antibiotic-susceptible *S*. Typhi ISP2825. N, negative control containing all PCR components except for *S*. Typhi genomic DNA. See Table 3 for details.

### Molecular basis of fluoroquinolone-resistance among XDR *S*. Typhi isolates

The acquisition of an IncY region harboring *qnrS* and one or more point mutations on the genes *gyrA, gyrB, parC*, and/or *parE*, referred to as the quinolone resistance determining region (QRDR), have been correlated to fluoroquinolone-resistance among typhoidal *Salmonella* strains. A PCR primer set specific for *qnrS* was designed and used to investigate whether XDR *S*. Typhi isolates encode this fluoroquinolone-resistance-related gene. We found that, unlike antibiotic-susceptible *S*. Typhi ISP2825, all XDR *S*. Typhi isolates tested encode *qnrS* (Figure 3A). Fluoroquinolone-resistant *Salmonella* strains have been reported to carry point mutations in *gyrA, gyrB, parC*, and/or *parE*, which exhibits variations depending on geographical locations (10, 12-20) (Table 4). Specific primer sets for the known mutations on these four genes were designed for PCR and PCR amplicon sequencing (Table 3). Consistent with their essential roles in bacterial cell replication, both antibiotic-susceptible *S*. Typhi ISP2825 and all XDR *S*. Typhi isolates resulted in PCR products with expected size for the four topoisomerase genes (Figure 3B-E). To determine whether XDR *S*. Typhi isolates resistant to fluoroquinolones have point mutations in these topoisomerase genes, we carried out Sanger sequencing of PCR amplicons. Consistent with the antibiotic-susceptible phenotype, *S*. Typhi ISP2825 carries wild-type topoisomerases. In contrast, we found that all the XDR *S*. Typhi isolates carry a mutant form of *gyrA* encoding for GyrA^Ser83Phe^ (Figure 3F-G). GyrA^Ser83Phe^ has been found most commonly among XDR *S*. Typhi identified from other endemic regions. We also found that these XDR *S*. Typhi isolates from pediatric septicemia patients encode wild-type *gyrB, parC*, and *parE* (Figure 3G and Table 4).

**Figure 3.**
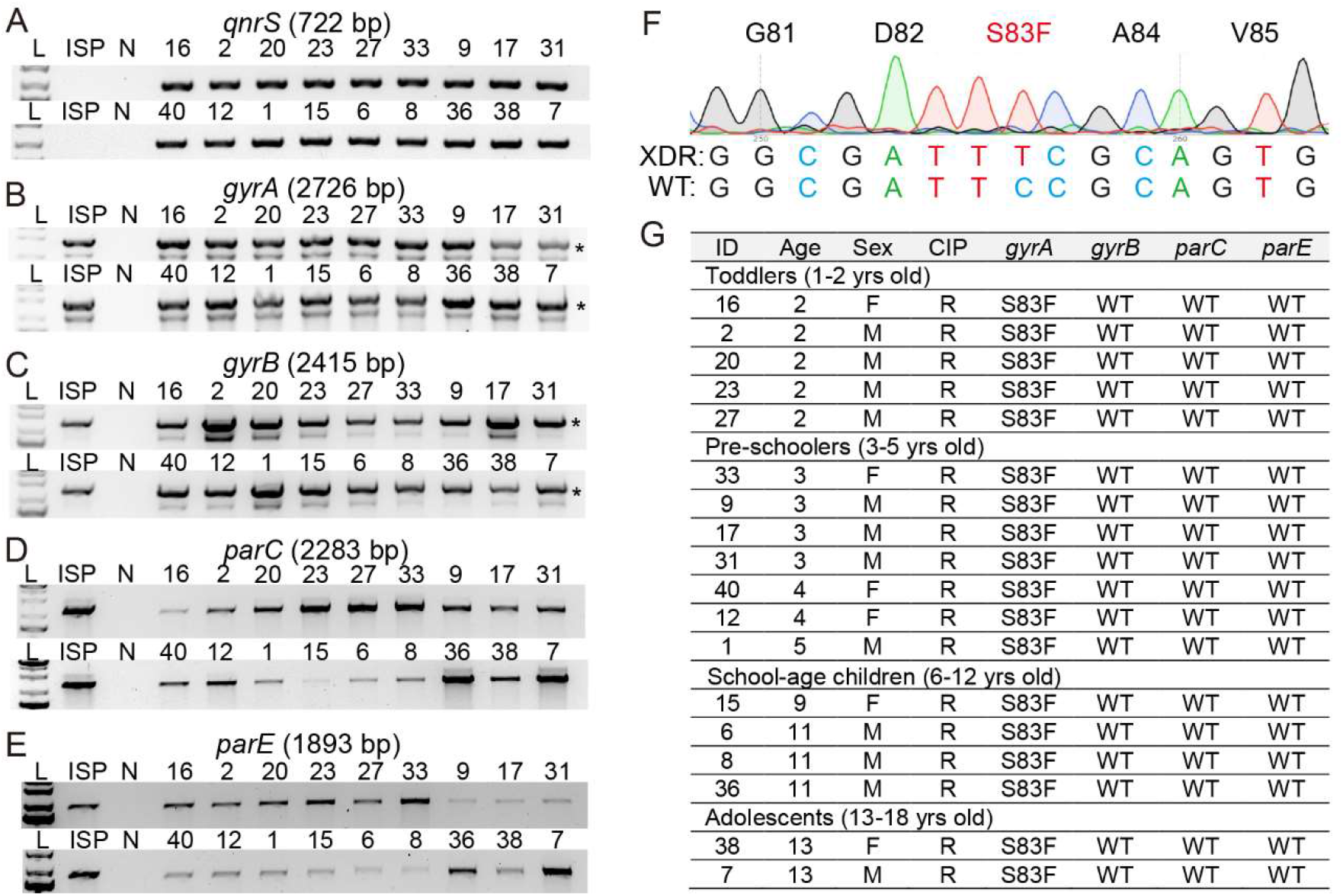
Molecular basis of fluoroquinolone-resistance among XDR *S*. Typhi isolates. **A-E**, PCR reactions for *qnrS* (**A**), *gyrA* (**B**), *gyrB* (**C**), *parC* (**D**), and *parE* (**E**). See Table 3 for details. ISP, antibiotic-susceptible *S*. Typhi ISP2825. N, negative control containing all PCR components except for *S*. Typhi genomic DNA. *, PCR amplicons specific to *gyrA* in B and *gyrB* in C. **F**, Representative sequencing chromatogram showing GyrA S83F mutation. **G**, Summary of PCR amplicon sequencing analysis for *gyrA, gyrB, parC*, and *parE*. See Table 4 for additional information. WT, wild type for the known mutations.

**Table 4.** Sequencing results associated with fluoroquinolone resistance, related to Fig. 3.

| Genes | Mutations | References | Findings |
| --- | --- | --- | --- |
| *gyrA* | M52L | (14) | Wild type |
|  | G81C | (16) | Wild type |
|  | D82G | (16) | Wild type |
|  | S83F/Y/L | (13, 20) | S83F |
|  | D87Y/H/N/G/A | (13, 20, 48) | Wild type |
|  | A119E | (16) | Wild type |
| *gyrB* | S464Y/F/T | (17) | Wild type |
|  | Q465L |  | Wild type |
|  | E466D |  | Wild type |
|  | A468E |  | Wild type |
| *parC* | T57S | (18, 19) | Wild type |
|  | G72S | (13) | Wild type |
|  | G78D | (16) | Wild type |
|  | D79G/R | (16, 18) | Wild type |
|  | S80R/I | (16, 20) | Wild type |
|  | G84G/K | (18, 20) | Wild type |
|  | W106G | (15) | Wild type |
| *parE* | D420N | (20) | Wild type |
|  | Y434S | (18) | Wild type |
|  | S458P | (16) | Wild type |

### Molecular basis of third-generation cephalosporin-resistance among XDR *S*. Typhi isolates

In addition to GyrA^Ser83Phe^, resistance to second-line antibiotics among XDR *S*. Typhi in Pakistan has been associated with an IncY region carrying the quinolone resistance gene *qnrS* (Figure 3A) and extended-spectrum ◸-lactamase resistance gene *blaCTX-M-15* (9, 38). Consistent with resistance to second-line antibiotics, fluoroquinolones and cephalosporins, among these XDR *S*. Typhi isolates from pediatric septicemia patients, we found the acquisition of *blaCTX-M-15* among all XDR *S*. Typhi tested (Figure 4). In contrast, antibiotic-susceptible *S*. Typhi ISP2825 did not result in PCR amplicon for *blaCTX-M-15*, indicating the specificity of the primers used.

**Figure 4.**
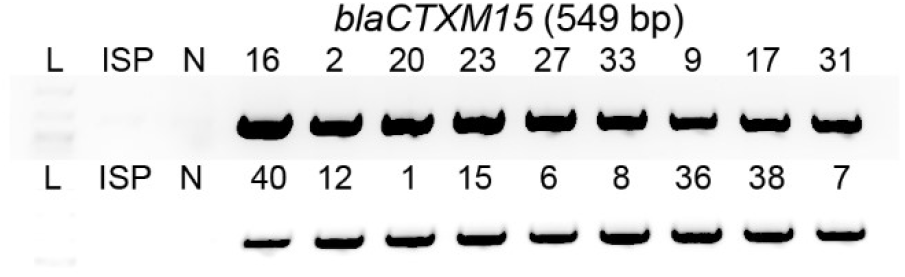
Molecular basis of third-generation cephalosporin-resistance among XDR *S*. Typhi isolates. PCR reactions for *blaCTXM15*. ISP, antibiotic-susceptible *S*. Typhi ISP2825. N, negative control containing all PCR components except for *S*. Typhi genomic DNA.

### Effects of efflux pumps on antibiotic resistance among XDR *S*. Typhi isolates

XDR *S*. Typhi isolates from pediatric typhoid patients exhibit some variations in their antibiotic-resistance/susceptibility profiles across antibiotics tested in Table 1. These results led us to investigate drug efflux pumps in XDR *S*. Typhi isolates since drug efflux pump systems are associated with resistance/susceptibility to a wide range of antibiotics. The recent worrisome trend among some XDR *S*. Typhi includes a correlation between efflux pump mutations and azithromycin resistance. For instance, a point mutation(s) on the antibiotic-binding subunit AcrB of the tripartite AcrAB-TolC efflux pumps (e.g., R717Q or R717L) has recently been correlated to azithromycin-resistance among some *S*. Typhi and *S*. Paratyphi A clinical isolates (19, 25, 26). In addition, *Salmonella* encodes another tripartite efflux pump, ABC-type MacAB-TolC, and three other small efflux pumps, major facilitator superfamily (MFS), multidrug and toxin extrusion (MATE), and small multidrug resistance (SMR), spanning in the inner membrane of the bacteria and therefore need to cooperate with another tripartite efflux pump such as RND-type AcrAB-TolC in exporting antibiotics (24). In *Neisseria gonorrhoeae* that, like *S*. Typhi, is also a human-adapted gram-negative bacterial pathogen, point mutations on the promoter of *macA* of the tripartite MacAB-TolC efflux pump have been correlated to azithromycin-resistance (39).

To assess whether point mutations on tripartite efflux pumps have occurred among XDR *S*. Typhi isolates from pediatric patients, we have determined *macA* promoter and *acrB* sequences via PCR and PCR amplicon sequencing using specific primer sets summarized in Table 3. We found that all XDR *S*. Typhi isolates tested carry wild-type -10 promoter sequence in the *macA* promoter and wild-type Arg at position 717 on the AcrB protein (Figure 5A-C). These results are consistent with azithromycin-susceptibility (2-8 µg/ml) among XDR *S*. Typhi isolates from our pediatric septicemia patient cohort (Figure 5C and Table 1).

**Figure 5.**
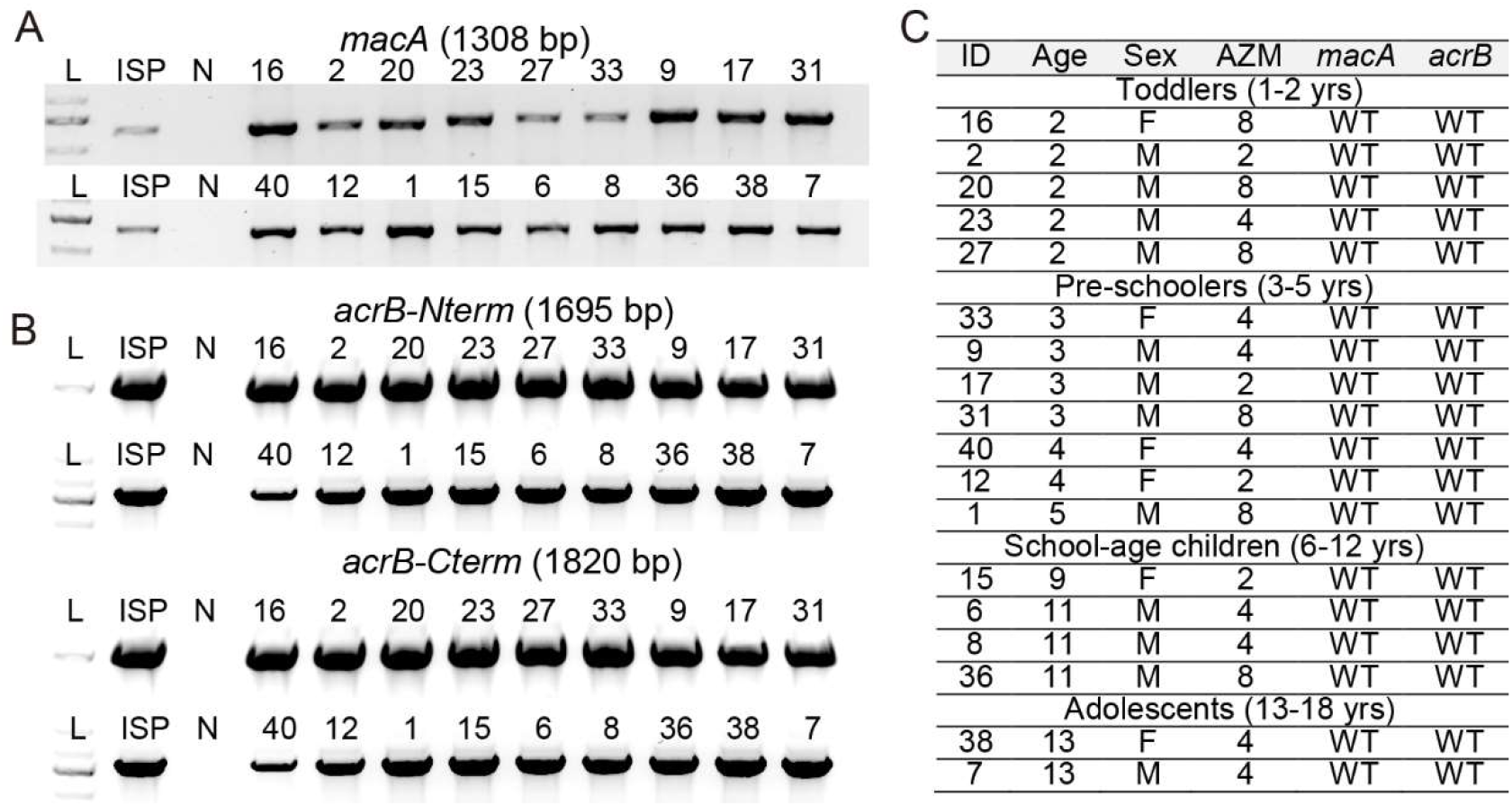
*macA* and *acrB* sequence analysis among XDR *S*. Typhi isolates. **A-B**, PCR reactions for *macA* (**A**) and *acrB* (**B**). *acrB* was split into two pieces for more productive PCR reaction outcomes (*acrB*-Nterm and *acrB*-Cterm). ISP, antibiotic-susceptible *S*. Typhi ISP2825. N, negative control containing all PCR components except for *S*. Typhi genomic DNA. See Table 3 for details. **C**, Summary of PCR amplicon sequencing analysis for *macA* and *acrB*. WT, wild type for the known mutations.

The expression of drug-efflux pumps is tightly regulated. For instance, AcrR represses *acrAB* gene expression by binding to the operator and inhibiting the transcription of *acrAB*. In 3-dimensional protein structure, wild-type AcrR protein monomer forms nine α-helices crucial for homodimer assembly, DNA-binding, and ligand-binding for its function in repressing *acrAB* expression (40, 41). Through whole-genome sequencing analysis of the latest NCBI RefSeq dataset of the fully assembled complete genome of *S*. Typhi (107 in total; Table 5), we found that two *S*. Typhi strains (RefSeq assembly accession IDs GCF_001121865.2 and GCF_900205275.1) carry a variant form of AcrR with 48 amino acid difference at the C-terminus (Figure 6A). Unlike the repressor AcrR variant, these two *S*. Typhi strains still carry wild-type RobA (NP_463442.1) and MarA (WP_000091194.1), activators for the *acrAB* gene expression, and wild-type AcrA (NP_459471.1), AcrB (NP_459470.1), and MacA (NP_459918.1) (data not shown). These results led us to investigate whether our XDR *S*. Typhi isolates carry a variant form of AcrR. Consistent with our azithromycin-susceptibility data, we found that all the XDR *S*. Typhi isolates from our pediatric septicemia patient cohort carry wild-type AcrR (Figure 6B-C).

**Table 5.** Details of the 107 completed *S*. Typhi genomes used in the study, related to Fig. 6.

| Accession number | Assembly number | BioProject | BioSample | BioSample ID | Serovar |
| --- | --- | --- | --- | --- | --- |
| GCF_000007545.1 | ASM754v1 | PRJNA371 | SAMN02604095 | 2604095 | Typhi |
| GCF_000195995.1 | ASM19599v1 | PRJNA236 | SAMEA1705914 | 25445 | Typhi |
| GCF_000245535.1 | ASM24553v1 | PRJNA80939 | SAMN02603101 | 2603101 | Typhi |
| GCF_000385905.1 | ASM38590v1 | PRJNA34855 | SAMN02603210 | 2603210 | Typhi |
| GCF_001048035.2 | ERL103914 | PRJEB3215 | SAMEA2072815 | 2363281 | Typhi |
| GCF_001048375.2 | M223 | PRJEB3215 | SAMEA2156512 | 2372886 | Typhi |
| GCF_001095585.2 | ERL114000 | PRJEB3215 | SAMEA2072817 | 2363290 | Typhi |
| GCF_001104165.2 | E00-7866 | PRJEB3215 | SAMEA2072798 | 2363309 | Typhi |
| GCF_001104885.2 | 10349_1#89_2 | PRJEB3215 | SAMEA2072799 | 2363310 | Typhi |
| GCF_001118185.2 | H12ESR00394-001A | PRJEB3215 | SAMEA2150110 | 2379332 | Typhi |
| GCF_001119245.2 | 76_1292 | PRJEB3215 | SAMEA1930246 | 2387556 | Typhi |
| GCF_001121865.2 | 404Ty | PRJEB3215 | SAMEA2072936 | 2363311 | Typhi |
| GCF_001127485.2 | ERL041834 | PRJEB3215 | SAMEA2072498 | 2363237 | Typhi |
| GCF_001135805.2 | ERL072973 | PRJEB3215 | SAMEA2072503 | 2363252 | Typhi |
| GCF_001148125.2 | ERL024120 | PRJEB3215 | SAMEA2072494 | 2363225 | Typhi |
| GCF_001148305.2 | ERL082356 | PRJEB3215 | SAMEA2072647 | 2363255 | Typhi |
| GCF_001163025.2 | H12ESR04734-001A | PRJEB3215 | SAMEA2072794 | 2363303 | Typhi |
| GCF_001165785.2 | ERL024919 | PRJEB3215 | SAMEA2072495 | 2363228 | Typhi |
| GCF_001302605.1 | ASM130260v1 | PRJNA286155 | SAMN03765654 | 3765654 | Typhi |
| GCF_001357935.2 | 80_2002 | PRJEB3215 | SAMEA1930249 | 2387559 | Typhi |
| GCF_001360555.2 | 2010_7898 | PRJEB3215 | SAMEA2058419 | 2301303 | Typhi |
| GCF_001362095.2 | 034151_4 | PRJEB3215 | SAMEA2072676 | 2363233 | Typhi |
| GCF_001362135.2 | 11909_3 | PRJEB3215 | SAMEA2072788 | 2363294 | Typhi |
| GCF_001362195.2 | H12ESR00755-001A | PRJEB3215 | SAMEA2072793 | 2363301 | Typhi |
| GCF_001362335.2 | Ty2 | PRJEB3215 | SAMEA2072934 | 2363305 | Typhi |
| GCF_003429465.1 | ASM342946v1 | PRJNA398278 | SAMN07507058 | 7507058 | Typhi |
| GCF_003716995.1 | ASM371699v1 | PRJNA474465 | SAMN09320528 | 9320528 | Typhi |
| GCF_003717015.1 | ASM371701v1 | PRJNA474465 | SAMN09320527 | 9320527 | Typhi |
| GCF_003717035.1 | ASM371703v1 | PRJNA474465 | SAMN09320526 | 9320526 | Typhi |
| GCF_003717055.1 | ASM371705v1 | PRJNA474465 | SAMN09320523 | 9320523 | Typhi |
| GCF_003717075.1 | ASM371707v1 | PRJNA474465 | SAMN09320522 | 9320522 | Typhi |
| GCF_003717095.1 | ASM371709v1 | PRJNA474465 | SAMN09320521 | 9320521 | Typhi |
| GCF_003717115.1 | ASM371711v1 | PRJNA474465 | SAMN09320520 | 9320520 | Typhi |
| GCF_003717135.1 | ASM371713v1 | PRJNA474465 | SAMN09320519 | 9320519 | Typhi |
| GCF_003717215.1 | ASM371721v1 | PRJNA474465 | SAMN09320518 | 9320518 | Typhi |
| GCF_003717285.1 | ASM371728v1 | PRJNA474465 | SAMN09320517 | 9320517 | Typhi |
| GCF_003717355.1 | ASM371735v1 | PRJNA474465 | SAMN09320516 | 9320516 | Typhi |
| GCF_003717395.1 | ASM371739v1 | PRJNA474465 | SAMN09320514 | 9320514 | Typhi |
| GCF_003717435.1 | ASM371743v1 | PRJNA474465 | SAMN09320513 | 9320513 | Typhi |
| GCF_003717455.1 | ASM371745v1 | PRJNA474465 | SAMN09320512 | 9320512 | Typhi |
| GCF_003717475.1 | ASM371747v1 | PRJNA474465 | SAMN09320511 | 9320511 | Typhi |
| GCF_003717515.1 | ASM371751v1 | PRJNA474465 | SAMN09320510 | 9320510 | Typhi |
| GCF_003717535.1 | ASM371753v1 | PRJNA474465 | SAMN09320509 | 9320509 | Typhi |
| GCF_003717575.1 | ASM371757v1 | PRJNA474465 | SAMN09320507 | 9320507 | Typhi |
| GCF_003717615.1 | ASM371761v1 | PRJNA474465 | SAMN09320564 | 9320564 | Typhi |
| GCF_003717635.1 | ASM371763v1 | PRJNA474465 | SAMN09320563 | 9320563 | Typhi |
| GCF_003717655.1 | ASM371765v1 | PRJNA474465 | SAMN09320562 | 9320562 | Typhi |
| GCF_003717675.1 | ASM371767v1 | PRJNA474465 | SAMN09320561 | 9320561 | Typhi |
| GCF_003717695.1 | ASM371769v1 | PRJNA474465 | SAMN09320560 | 9320560 | Typhi |
| GCF_003717715.1 | ASM371771v1 | PRJNA474465 | SAMN09320559 | 9320559 | Typhi |
| GCF_003717735.1 | ASM371773v1 | PRJNA474465 | SAMN09320558 | 9320558 | Typhi |
| GCF_003717755.1 | ASM371775v1 | PRJNA474465 | SAMN09320557 | 9320557 | Typhi |
| GCF_003717775.1 | ASM371777v1 | PRJNA474465 | SAMN09320556 | 9320556 | Typhi |
| GCF_003717795.1 | ASM371779v1 | PRJNA474465 | SAMN09320555 | 9320555 | Typhi |
| GCF_003717815.1 | ASM371781v1 | PRJNA474465 | SAMN09320554 | 9320554 | Typhi |
| GCF_003717835.1 | ASM371783v1 | PRJNA474465 | SAMN09320553 | 9320553 | Typhi |
| GCF_003717855.1 | ASM371785v1 | PRJNA474465 | SAMN09320552 | 9320552 | Typhi |
| GCF_003717875.1 | ASM371787v1 | PRJNA474465 | SAMN09320551 | 9320551 | Typhi |
| GCF_003717895.1 | ASM371789v1 | PRJNA474465 | SAMN09320550 | 9320550 | Typhi |
| GCF_003717915.1 | ASM371791v1 | PRJNA474465 | SAMN09320549 | 9320549 | Typhi |
| GCF_003717935.1 | ASM371793v1 | PRJNA474465 | SAMN09320548 | 9320548 | Typhi |
| GCF_003717955.1 | ASM371795v1 | PRJNA474465 | SAMN09320547 | 9320547 | Typhi |
| GCF_003717975.1 | ASM371797v1 | PRJNA474465 | SAMN09320546 | 9320546 | Typhi |
| GCF_003717995.1 | ASM371799v1 | PRJNA474465 | SAMN09320545 | 9320545 | Typhi |
| GCF_003718015.1 | ASM371801v1 | PRJNA474465 | SAMN09320544 | 9320544 | Typhi |
| GCF_003718035.1 | ASM371803v1 | PRJNA474465 | SAMN09320543 | 9320543 | Typhi |
| GCF_003718055.1 | ASM371805v1 | PRJNA474465 | SAMN09320542 | 9320542 | Typhi |
| GCF_003718075.1 | ASM371807v1 | PRJNA474465 | SAMN09320541 | 9320541 | Typhi |
| GCF_003718095.1 | ASM371809v1 | PRJNA474465 | SAMN09320540 | 9320540 | Typhi |
| GCF_003718115.1 | ASM371811v1 | PRJNA474465 | SAMN09320539 | 9320539 | Typhi |
| GCF_003718135.1 | ASM371813v1 | PRJNA474465 | SAMN09320538 | 9320538 | Typhi |
| GCF_003718155.1 | ASM371815v1 | PRJNA474465 | SAMN09320537 | 9320537 | Typhi |
| GCF_003718175.1 | ASM371817v1 | PRJNA474465 | SAMN09320536 | 9320536 | Typhi |
| GCF_003718195.1 | ASM371819v1 | PRJNA474465 | SAMN09320535 | 9320535 | Typhi |
| GCF_003718235.1 | ASM371823v1 | PRJNA474465 | SAMN09320533 | 9320533 | Typhi |
| GCF_003718255.1 | ASM371825v1 | PRJNA474465 | SAMN09320532 | 9320532 | Typhi |
| GCF_003718275.1 | ASM371827v1 | PRJNA474465 | SAMN09320531 | 9320531 | Typhi |
| GCF_003718295.1 | ASM371829v1 | PRJNA474465 | SAMN09320530 | 9320530 | Typhi |
| GCF_003718315.1 | ASM371831v1 | PRJNA474465 | SAMN09320529 | 9320529 | Typhi |
| GCF_003718355.1 | ASM371835v1 | PRJNA474465 | SAMN09320578 | 9320578 | Typhi |
| GCF_003718375.1 | ASM371837v1 | PRJNA474465 | SAMN09320577 | 9320577 | Typhi |
| GCF_003718395.1 | ASM371839v1 | PRJNA474465 | SAMN09320576 | 9320576 | Typhi |
| GCF_003718415.1 | ASM371841v1 | PRJNA474465 | SAMN09320575 | 9320575 | Typhi |
| GCF_003718435.1 | ASM371843v1 | PRJNA474465 | SAMN09320574 | 9320574 | Typhi |
| GCF_003718455.1 | ASM371845v1 | PRJNA474465 | SAMN09320573 | 9320573 | Typhi |
| GCF_003718475.1 | ASM371847v1 | PRJNA474465 | SAMN09320572 | 9320572 | Typhi |
| GCF_003718495.1 | ASM371849v1 | PRJNA474465 | SAMN09320571 | 9320571 | Typhi |
| GCF_003718515.1 | ASM371851v1 | PRJNA474465 | SAMN09320570 | 9320570 | Typhi |
| GCF_003718535.1 | ASM371853v1 | PRJNA474465 | SAMN09320569 | 9320569 | Typhi |
| GCF_003718555.1 | ASM371855v1 | PRJNA474465 | SAMN09320568 | 9320568 | Typhi |
| GCF_003718575.1 | ASM371857v1 | PRJNA474465 | SAMN09320567 | 9320567 | Typhi |
| GCF_003718595.1 | ASM371859v1 | PRJNA474465 | SAMN09320566 | 9320566 | Typhi |
| GCF_003718615.1 | ASM371861v1 | PRJNA474465 | SAMN09320565 | 9320565 | Typhi |
| GCF_003718635.1 | ASM371863v1 | PRJNA474465 | SAMN09320525 | 9320525 | Typhi |
| GCF_003718655.1 | ASM371865v1 | PRJNA474465 | SAMN09320506 | 9320506 | Typhi |
| GCF_003719215.1 | ASM371921v1 | PRJNA474465 | SAMN09320534 | 9320534 | Typhi |
| GCF_003719235.1 | ASM371923v1 | PRJNA474465 | SAMN09320508 | 9320508 | Typhi |
| GCF_003719255.1 | ASM371925v1 | PRJNA474465 | SAMN09320515 | 9320515 | Typhi |
| GCF_003719555.1 | ASM371955v1 | PRJNA471337 | SAMN09208111 | 9208111 | Typhi |
| GCF_004136335.1 | ASM413633v1 | PRJNA480202 | SAMN09630442 | 9630442 | Typhi |
| GCF_005885835.1 | ASM588583v1 | PRJNA543969 | SAMN11792777 | 11792777 | Typhi |
| GCF_900185485.1 | BL60006 | PRJEB21155 | SAMEA1041091<br>93 | 7190870 | Typhi |
| GCF_900205255.1 | 1554 | PRJEB5919 | SAMEA3109638 | 3338098 | Typhi |
| GCF_900205265.1 | lupe_GEN005<br>9_5 | PRJEB5919 | SAMEA2564024 | 3071773 | Typhi |
| GCF_900205275.1 | 403Ty | PRJEB5919 | SAMEA2564027 | 3071775 | Typhi |
| GCF_900205295.1 | E98-3139 | PRJEB5919 | SAMEA2467787 | 3000319 | Typhi |
| GCF_901457615.1 | ERS3381924 | PRJEB32272 | SAMEA5577690 | 11516613 | Typhi |

**Figure 6.**
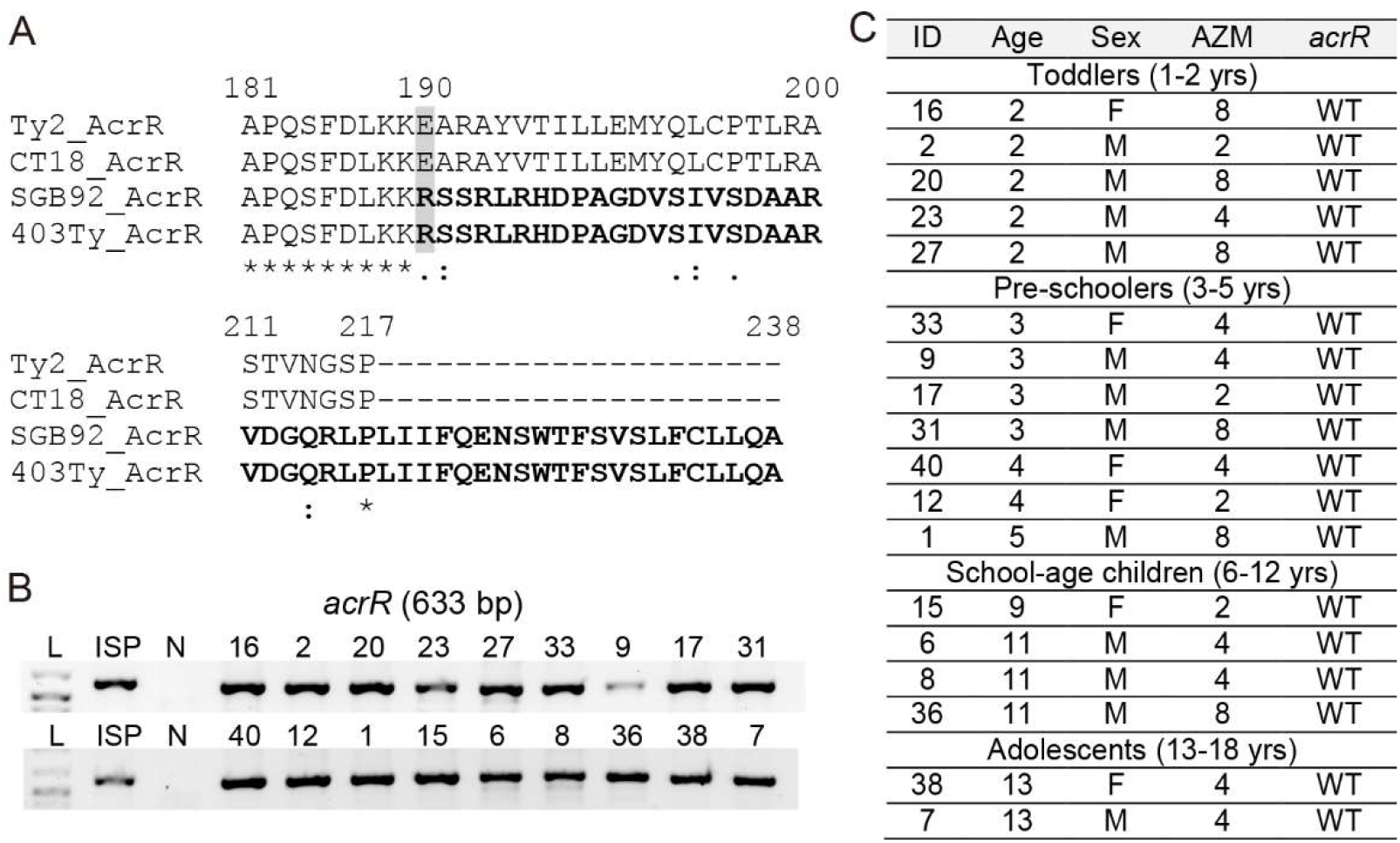
AcrR sequence analysis among XDR *S*. Typhi isolates. **A**, AcrR amino acid sequence comparison analysis of the latest RefSeq dataset of the fully assembled complete genome of *S*. Typhi (107 in total) collected from NCBI as of Feb 26, 2021. Ty2, *S*. Typhi Ty2 (RefSeq assembly accession: GCF_000007545.1, assembly name: ASM754v1, strain: Ty2, submitter: University of Wisconsin). CT18, *S*. Typhi CT18 (RefSeq assembly accession: GCF_000195995.1, assembly name: ASM19599v1, strain: CT18, submitter: Sanger Institute). SGB92, *S*. Typhi SGB92 (RefSeq assembly accession: GCF_001121865.2, assembly name: 404Ty, strain name: SGB92, submitter: Wellcome Sanger Institute). 403Ty, *S*. Typhi 403Ty-sc-1979084 (RefSeq assembly accession: GCF_900205275.1, assembly name: 403Ty, isolate: 403Ty-sc-1979084, submitter: Wellcome Sanger Institute). See Table 5 for details. **B**, PCR reactions for *acrR*. ISP, antibiotic-susceptible *S*. Typhi ISP2825. N, negative control containing all PCR components except for *S*. Typhi genomic DNA. **C**, Summary of PCR amplicon sequencing analysis for *acrR*. WT, wild type for the known mutations.

## Discussion

XDR *S*. Typhi is more common among pediatric patients but the majority of antibiotic resistance studies available have been carried out using *S*. Typhi isolates from adult patients. Here, we characterized *S*. Typhi isolates from a medium size cohort of pediatric typhoid patients to determine antibiotic-resistance-related gene signatures associated with their drug-resistant profiles. This study provides a valuable overview of the recent (2019-2020) populations in the setting of Lahore, Pakistan, among a septic pediatric cohort and provides insights into the development of simple, cost-effective molecular detection methods with point-of-care testing potential.

Biochemical method-mediated identification assisted by the streamlined automated system was further validated via molecular typing for typhoidal *Salmonella* specific gene sequences (42-44). Antibiotic resistance profiles were obtained through the automated test system that followed the CLSI 2018 guidelines. To determine the molecular basis of the antibiotic resistance profiles among these XDR *S*. Typhi, we have designed primer sets and optimized PCR reaction conditions for antibiotic resistance genes harbored in the IncHI1 (*catA1, blaTEM1, dhfR7*, and *sul1*) and IncY (*qnrS and blaCTX-M-15*) regions, contributing to resistance to front-line and second-line antibiotics, respectively. Overall findings across the XDR *S*. Typhi isolates for these antibiotic resistance genes are in agreement with our MIC results and other reports correlated to molecular determinants of MDR and XDR phenotypes among recent XDR *S*. Typhi isolates from adult typhoid patients (9).

Besides *qnrS*, mutations on *gyrA, gyrB, parC*, and *parE* also contribute to fluoroquinolone-resistance, which exhibits more diverse patterns depending on geographical locations. Our XDR *S*. Typhi isolates from pediatric patients in Northern Pakistan carry GyrA^S83F^ across all samples tested, while the remaining genes were found to be wild-type (Figure 3 and Table 4). This result indicates that XDR *S*. Typhi circulating in this geographical location is different from the ones prevalent in other locations that exhibit different QRDR mutation signatures. Future investigations on XDR *S*. Typhi isolates from adult patients in the same geographical location would inform us about whether a divergent host adaptation process has occurred in pediatric and adult patients.

Macrolides such as azithromycin are considered the only remaining oral antibiotic option in treating XDR *S*. Typhi resistant to both front-line and second-line antibiotics. An additional treatment option against XDR *S*. Typhi, although requiring injection, are carbapenems such as imipenem and meropenem. Our XDR *S*. Typhi isolates from pediatric patients are susceptible to azithromycin, imipenem, and meropenem (Table 1). In contrast, 5% and 48% of recent *S*. Typhi isolates (n=81) from a mid-size adult cohort in Northern Punjab were resistant to azithromycin and meropenem, respectively (36), supporting the concept of a divergent host adaptation and/or transmission process in child and adult groups. These results also support a possibility that, with time, molecular determinants for azithromycin-resistance and meropenem-resistance would likely be adopted in nearly all *S*. Typhi circulating locally and globally.

Current analysis indicates that our XDR *S*. Typhi isolates from pediatric septicemia patients in Punjab do not carry molecular determinants that have been correlated to azithromycin-resistance in typhoidal *Salmonella, S*. Typhi and *S*. Paratyphi A, and another human-adapted Gram-negative pathogen *N. gonorrhoeae* (Figure 5). Besides wild-type AcrB and wild-type -10 promoter sequence in the *macA* promoter across our XDR *S*. Typhi isolates, WGS analysis of the latest NCBI RefSeq dataset of the fully assembled complete genome of *S*. Typhi (107 in total), conducted as part of this study, indicates the emergence of *S*. Typhi strains carrying the frameshifted AcrR variant, the repressor of the *acrAB* efflux pump components.

Although further investigations are required to understand the consequence of having the AcrR frameshifted variant in antibiotic-resistance, it is intriguing to hypothesize that the AcrR variant is less effective in repressing the expression of *acrAB*, therefore contributing to antibiotic-resistance such as azithromycin. We also found that those *S*. Typhi strains carrying the AcrR variant carry wild-type RobA and MarA, activators for the *acrAB* gene expression, and wild-type AcrAB, collectively supporting the hypothesis that possession of the AcrR frameshifted variant is an adaptation/evolution outcome, rather than a stochastic event outcome.

The emergence and spread of *S*. Typhi resistant to macrolides and carbapenems are a serious global health concern, deserving close surveillance for local and global spread. We envision that some of the methods detecting key molecular determinants for *S*. Typhi antibiotic resistance used in the current study could be developed as a surveillance strategy and point-of-care testing strategy. The detection and analysis methods for resistance to front-line and second-line antibiotics described in the study are straightforward. Besides efflux pump related molecular determinants described in the study, the future surveillance strategy could include additional molecular traits predicted to be associated with resistance to macrolides and carbapenems in *S*. Typhi. For instance, in *Enterobacteriaceae, erm* genes encoding for methylases to modify target sites, *ere* genes for esterase transferases, and *mph* genes for phosphor transferases are known to confer macrolide-resistance by altering the structure of antibiotics (45). Similarly, carbapenem resistance in *S*. Typhi can be acquired by mutational events or gene acquisition via horizontal gene transfer, leading to the overexpression of efflux pumps that expel carbapenems and the acquisition of carbapenemases. The most effective carbapenemases known that hydrolyze carbapenem and spread across many bacterial pathogens are KPC, VIM, IMP, NDM and OXA-48 types (46).

In summary, this study informs the molecular basis of antibiotic-resistance among recent *S*. Typhi isolates from pediatric septicemia patients and provides insights into the development of molecular detection and control strategies for XDR *S*. Typhi.

## Materials & Methods

### Ethics Statement

Before initiating this research, ethical approval was obtained following the Declaration of Helsinki from the Institutional Review Board (IRB# FMH-03-2020-IRB-774-F), the Fatima Memorial Hospital Lahore. In addition, informed consent was obtained from a legal guardian of each study participant. Informed consent was read to the person in the language they understood and signed appropriately. They were willing to provide a sample and utilize the isolates for research. They were assured that the samples would be used solely for research purposes and that personal information would be kept confidential. Before samples were transferred to researchers, all XDR *S*. Typhi samples were de-identified, number-based identification codes were assigned to samples (Tables 1 and 2). The data were analyzed anonymously throughout the study.

### *S*. Typhi isolation from patient specimens and MIC determination

The BACT/ALERT® 3D Microbial Detection System with PF/PF plus culture bottles (bioMérieux, France), an automated bacterial culture and antibiotic-resistance test system capable of incubating, agitating, and continuously monitoring aerobic and anaerobic media inoculated with patient specimens was used in this study. The samples were collected between October 2019 to January 2020. In brief, 1-4 mL blood samples of each septicemia suspected child based on their age and bodyweight were taken and placed in BacT/ALERT PF/PF plus bottles for up to 5 days. The bottles contained BacT/Alert FAN Plus media with Adsorbent Polymeric Beads (APB) that neutralized antimicrobials (47). Positive blood culture bottles were sub-cultured on blood and MacConkey agar plates and incubated overnight at 37°C aerobically. Preliminary identification of the isolates was conducted according to colony morphology and culture characteristics. MIC determinations were made using the automatic VITEK 2 compact system (bioMérieux, France) and antibiotics interpretation was carried out as per the clinical laboratory standards institute (CLSI) 2018 guidelines (https://clsi.org).

### XDR *S*. Typhi samples selection for detailed molecular characterization

Of 45 XDR *S*. Typhi isolates, 18 isolates were selected for detailed molecular characterization based on their MIC results, gender, age, and hospital wards, representing all 45 XDR isolates (Tables 1 and 2).

### PCR-based detection of antibiotic-resistance-related genes among XDR *S*. Typhi isolates

Bacterial genomic DNA was prepared using a DNeasy bacterial DNA extraction kit (QIAGEN) following the vendor’s recommendation. The PCR primer sequences and reaction conditions used are summarized in Table 3. Green Taq DNA polymerase with provided buffers (GenScript) was used for *pltB, blaTEM1, dhfR7, sul1, catA1, parC, parE, blaCTXM15, macA, acrB*, and *acrR*. Phusion high fidelity DNA polymerase with the provided GC buffer (New England BioLabs) or Herculase II fusion DNA polymerase (Agilent) was used for *gyrA, gyrB*, and *qnrS*. PCR reaction steps were: pre-denaturation at 95°C for 3min, 34 cycles of denaturation at 95°C for 30 sec, annealing (see Table 3), and extension at 72°C for 1 kb/min (see Table 3 for amplicon size), and final extension at 72°C for 7 min using a C1000 Touch Thermal Cycle (BIO-RAD). PCR results were run on 1% agarose gels and imaged using an iBright CL1500 Imaging system (ThermoFisher Scientific).

### Sanger sequencing of PCR amplicons

When indicated, PCR amplicons were extracted from agarose gels for sequencing analysis by using the QIAEX Ⅱ gel extraction system (QIAGEN, cat # 20051), followed by standard Sanger sequencing (The Cornell Institute of Biotechnology or Eton Bioscience Inc). The primer sequences used for Sanger sequencing are summarized in Table 3.

### Whole-genome sequencing analysis for efflux pump-related genes

The latest NCBI RefSeq dataset of the fully assembled complete genome of *S*. Typhi (107 in total; Table 5) was collected on Feb 26, 2021. The 107 complete whole-genome sequences were utilized to analyze the sequence variations for *acrR* (NP_459472.1) using ’General Feature Formats (gff)’ files with a bash script (grep acrR *.gff | grep pseudo=true). Two whole genome sequences (GCF_001121865.2 and GCF_900205275.1) that have an *acrR* variant were further analyzed with CLC Main Workbench 8.1.3 (QIAGEN) for multidrug efflux pump-related genes: *macA* (NP_459918.1), *acrA* (NP_459471.1), *acrB* (NP_459470.1), *marA* (WP_000091194.1), and *robA* (NP_463442.1).

## Acknowledgments

This work was supported in part by NIH R01 AI137345, AI139625, and AI141514 to JS. The funders had no role in the study design, data collection and analysis, decision to publish, or preparation of the manuscript.

## Author contributions

C.K. conducted experiments shown in Figs. 1-6 using genomic DNA received from M.U.Q., and prepared the manuscript draft. D.P.N. contributed to experiments shown in Figs. 1-6. G.Y.L. conducted WGS analysis for efflux pump-related genes shown in Fig. 6. R.S.K. contributed to antibiotic resistance-related gene sequence analysis. I.L. A.B. and Q.A. isolated XDR *S*. Typhi strains, conducted MIC determination experiments, and genomic DNA preparations. M.U.Q supervised the isolation and antibiotic resistance phenotype characterization study. J.S. supervised the molecular characterization study using genomic DNA received, and prepared the manuscript with input from all authors.

